# Extracellular chaperones modulate AL light chains fibrillar aggregation and contribute to amyloid structural heterogeneity

**DOI:** 10.64898/2026.05.29.728721

**Authors:** Loredana Marchese, Mariagrazia Battaglia, P. Patrizia Mangione, Annalisa Relini, Giulia Codroico, Sara Raimondi, Federico Forneris, Silvia Faravelli, Beatrice Leonardini, Claudio Canale, Guglielmo Verona, Diana Canetti, Vittorio Bellotti, Sofia Giorgetti, Alessandra Corazza, Francesca Lavatelli

**Affiliations:** Pathology Unit, Fondazione IRCCS Policlinico San Matteo, Pavia, Italy; Department of Molecular Medicine, University of Pavia, Italy; Research Area, Fondazione IRCCS Policlinico San Matteo, Pavia, Italy; Department of Physics, University of Genoa, Italy; Department of Biology and Biotechnology, University of Pavia, Italy; UCL London, UK; Department of Medicine, University of Udine, Italy

## Abstract

In AL amyloidosis, monoclonal immunoglobulin light chains (LCs) aggregate as amyloid fibrils in tissues. In synergy with the intrinsic aggregation propensity of specific LC sequences, microenvironment factors may be involved in tuning the disease pathophysiology, in particular proteolytic LC remodelling and heterotypic interactions in the extracellular milieu. Accounting for extrinsic modulators is critical for understanding the phenotypic variability of AL, usually imputed mainly to the LC diversity. We investigated the effects of apolipoprotein E (allele 3, apoE3) and clusterin (CLU), two amyloid-signature proteins involved in extracellular proteostasis, on the fibrillogenesis kinetics of amyloidogenic LC fragments from patient-derived sequences, as well as on aggregate composition, fibril morphology and thermodynamic stability. We show that apoE3 and CLU act as heterotypic interactors of prefibrillar and fibrillar LCs, significantly modulating LC amyloidogenesis, with complex and non-monotypic effects that range from anti- to pro-amyloidogenic depending on their concentration and on the LC’s intrinsic amyloidogenicity. ApoE3 and CLU also influence fibril morphology, possibly by modifying protofilament association, and alter their thermodynamic properties. LC interactors may play a significant and insofar underappreciated role in the AL pathophysiology *in vivo*, likely contributing to phenotypic variability and structural polymorphisms and, possibly, to fibril resilience to amyloid reabsorption strategies.

## INTRODUCTION

Aggregation of immunoglobulin light chains (LCs) as extracellular fibrils is the hallmark of AL amyloidosis, a life-threatening disease associated with plasma or B cell clones that overproduce misfolding-prone monoclonal components (1). Progressive amyloid infiltration and concomitant proteotoxicity of prefibrillar LCs severely compromise the function of affected organs. Heart and kidney are most frequently involved, but virtually every extracerebral tissue can be affected. Compared to other amyloidoses, AL is characterized by heterogeneous clinical and pathological phenotypes, each patient presenting with a specific amyloid deposition pattern and disease severity. This variability is paralleled by the molecular diverseness of LCs, downstream products of the immunoglobulin maturation process with unique primary sequences, especially within the complementarity-determining regions (CDRs) of the variable domain (V_L_) (2).

Not all clonal LCs are associated with amyloid deposition *in vivo*; the repertoire of AL-causing LCs is skewed towards the λ isotype and preferential usage of specific germlines genes (2, 3). Several lines of evidence, however, indicate that the primary structure alone is not sufficient to explain LC fibril conversion and imply the pathogenetic contribution of additional events taking place in the microenvironment (4, 5). Amyloidogenesis occurs in the extracellular space of target organs, where exogenous factors may trigger aggregation and modulate fibril stability. The established evidence that *ex vivo* AL deposits are composed of a population of LC fragments missing part of the constant domain (C_L_) (5-7) and that the V_L_ alone forms the structured fibrillar core (8, 9) suggest that proteolytic remodelling, in first place, is required to unleash a LC’s amyloidogenic potential (4, 10). *In vitro*, the isolated V_L_ or fragments encompassing the V_L_ and part of the C_L_ (henceforth indicated as V_L_-C_L_ fragments) efficiently convert into amyloid fibrils under physiological conditions, in contrast to full-length LCs, which are remarkably resistant (5).

Investigation of affected tissues, moreover, displays microenvironment features that yield additional clues on the pathophysiological mechanisms, in particular extracellular matrix (ECM) modifications and accumulation of a plethora of non-resident amyloid-associated proteins, among which apolipoproteins (e.g. apoE, apoAIV), serum amyloid P (SAMP), clusterin (CLU) and vitronectin (VTN) (11). Some of these, such as CLU and apoE, are known to participate in extracellular proteostasis, by binding misfolded polypeptides and working as holdases that direct altered clients to cellular-mediated degradation (12, 13). Their ubiquitous co-deposition with amyloid, however, raises questions on their role in the disease processes, as fibril buildup ultimately indicates a failure in proteostasis mechanisms, which are either insufficient or even permissive in the chain of events converting folded precursors into aggregates. The clinical evidence that gene variants of apoE and CLU are among the most prominent risk factors for late onset Alzheimer’s disease, of which brain Aβ amyloid deposits are a hallmark (14-16), supports a pathophysiological contribution of these chaperones to amyloid pathology. It is indeed becoming clear that heterotypic interactions between amyloidogenic proteins and environmental components can profoundly affect the fibrillogenesis kinetics and modulate fibril properties (17-19). Binding of Aβ to distinct partners, for example, was suggested to modulate cellular vulnerability and polymorphic bias in different brain areas (17, 20, 21). The known properties of CLU, one of the most extensively characterized extracellular chaperones, exemplify the complex role that these proteins can play in modulating amyloid vulnerability. In fact, whereas clusterin inhibits fibrillar and amorphous aggregation of clients such as Aβ when present at sufficient concentrations, it can have a paradox pro-amyloidogenic effect in the presence of a large molar excess of the amyloidogenic substrate, possibly through stabilization of transient on-pathway conformers (22, 23).

Despite the unquestionable interest of this topic, the role of heterotypic interactors in extracerebral amyloidoses, including AL, is understudied. Accounting for environmental modulators of LC aggregation could be critical for understanding the vulnerability, as well as the phenotypic and clinical heterogeneity of this disease. In this perspective, we have investigated the effects of the paradigmatic amyloid signature proteins CLU and apoE on the fibrillogenesis of two amyloidogenic LCs belonging to distinct germline genes families, IGLV6-57 and IGLV1-51. We show that both CLU and apoE3 significantly modulate LC aggregation kinetics, fibril morphology and stability in a dose-dependent but not monotypic way, exerting protective or permissive effects depending on their concentration and on the light chain’s sequence. Our results suggest that amyloid-associated chaperones may play a significant and insofar underappreciated role in the pathophysiology of AL amyloidosis *in vivo*, with important translational implications.

## RESULTS

### The V_L_ sequence modulates the fibrillogenesis kinetics of amyloidogenic LC fragments

We investigated the influence of sequence-related and extrinsic factors in the amyloid aggregation of two recombinant V_L_-C_L_ fragments from patient-derived LC sequences belonging to IGLV6-57 and IGLV1-51 germline gene families. These fragments, indicated respectively as 133-AL55 (reported in (5)) and 128-ALH7, encompass the entire V_L_ region and 22 residues of the constant one, extending up to β-strand βAC of the C_L_ (Figure S1 A). Consistently with what previously shown for 133-AL55, the 128-ALH7 LC fragment is highly amyloidogenic and entirely converts into amyloid fibrils under physiological conditions of pH, ionic strength and temperature (Figure 1). Compared to 133-AL55, 128-ALH7 has a significantly longer lag phase (Figures 1 and 2). The corresponding full-length LC ALH7 (6, 24) is, conversely, remarkably resistant to amyloid formation (data not shown), analogously to what previously observed for full-length AL55 (5, 10). To explore sequence features related to amyloidogenicity, the presence of amyloid-prone regions (APR) in both fragments was predicted *in silico* using multiple sequence-based tools (AmylPred2 with consensus of four methods, Aggrescan, ANuPP) (25-27). A consensus score was calculated based on the number of algorithms that identified each residue as belonging to an APR (Figure S1 B). Overall, whereas multiple APRs are predicted in both LCs, their number is higher in 133-AL55, and only in this LC a consensus is reached by all algorithms on two specific segments (17-21 and 31-36) (Figure S1 D).

**Figure 1.**
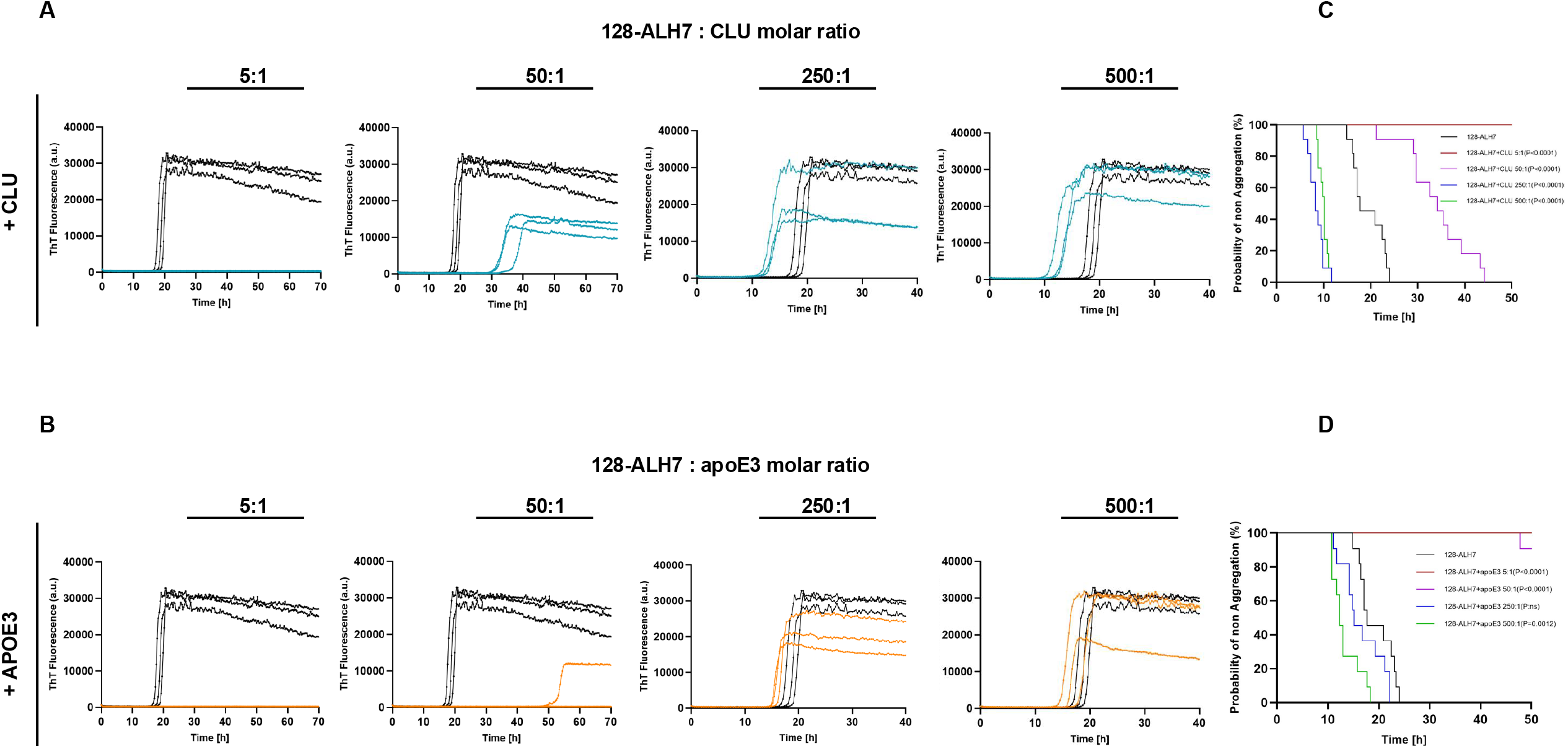
Fibrillogenesis kinetics of the amyloidogenic LC fragment 128-ALH7 in the presence of CLU and apoE3. Amyloid formation was monitored based on the increase in ThT fluorescence. 128-ALH7 (black curves in all panels) was used at a concentration of 10 μM in all tests, whereas CLU (panels in A, blue curves) and apoE3 (panels in B, orange curves) were added at substoichiometric concentrations ranging from 2 μM to 0.02 μM (i.e. 5:1 to 500:1 LC:cofactor molar ratios). Each curve in the panels refers to a single microplate well; data are representative of three independent experiments that included the full set of displayed conditions (3 to 5 wells/condition/experiment). The fibrillogenesis lag phase in the presence of various concentrations of CLU (panel C) and apoE3 (panel D) was evaluated using Kaplan-Meier analysis (n=11 in all conditions), considering as event the start time of fibril growth phase in the ThT plots. 128-ALH7 alone is displayed in black; colour codes for each condition are shown in the inset. Curves were compared using long-rank (Mantel-Cox) test with Bonferroni correction. In the case of CLU (panel C), all curves are significantly different from 128-ALH7 alone. In the case of apoE3 (panel D), all comparisons were significant except for 250:1 apoE3.

**Figure 2.**
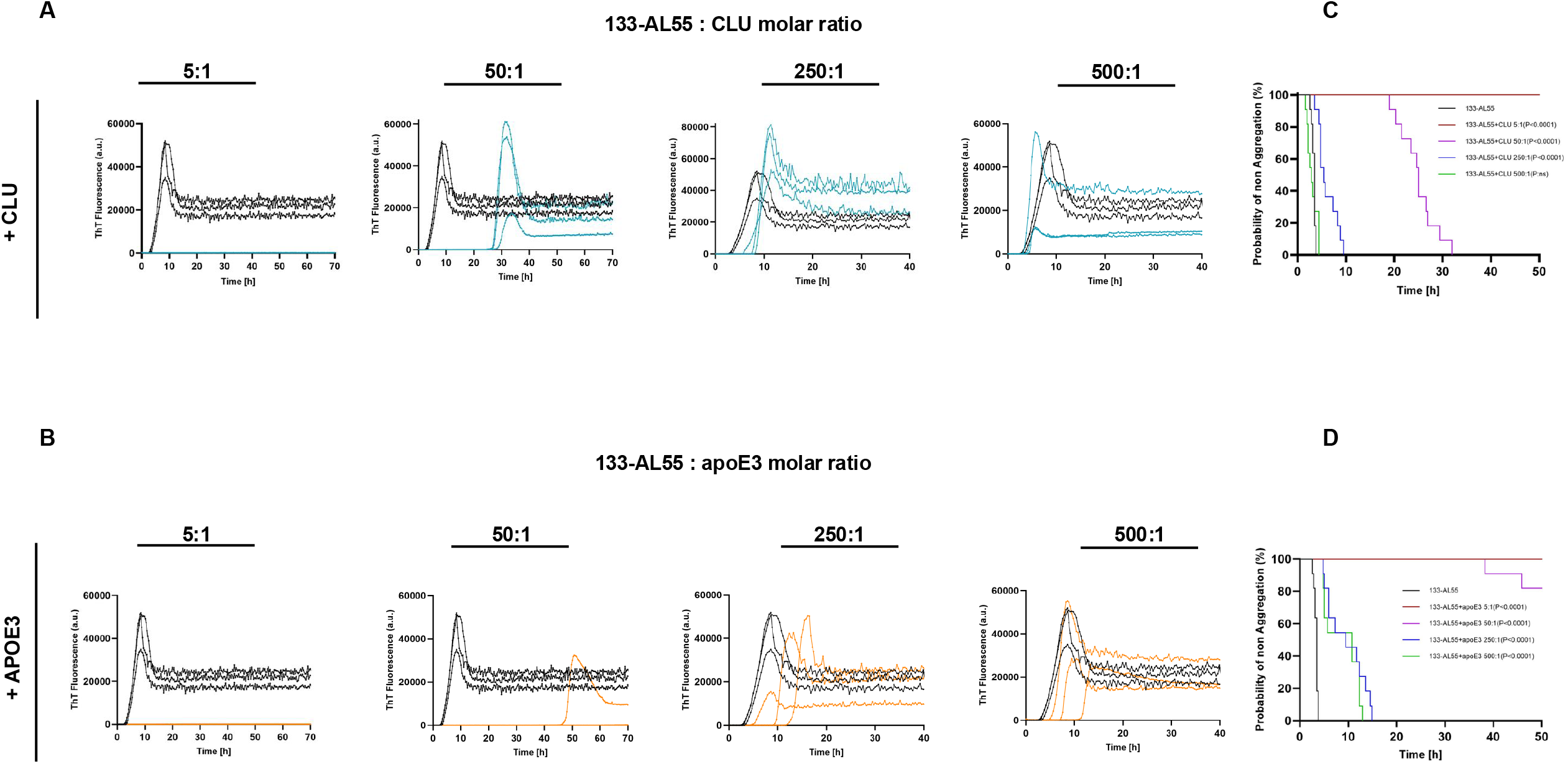
Fibrillogenesis kinetics of the amyloidogenic LC fragment 133-AL55 in the presence of CLU and apoE3. Amyloid formation was monitored based on the increase in ThT fluorescence. 133-AL55 (black curves in all panels) was used at a concentration of 10 μM in all tests, whereas CLU (panels in A, blue curves) and apoE3 (panels in B, orange curves) were added in substoichiometric concentrations ranging from 2 μM to 0.02 μM (i.e. 5:1 to 500:1 LCs:cofactors molar ratios). Each curve in the panels refers to a single microplate well; data are representative of three independent experiments including the full set of tested conditions. The fibrillogenesis lag phase in the presence of various concentrations of CLU (panel C) and ApoE3 (panel D) was evaluated using Kaplan-Meier analysis, as described in Figure 1 (n=11 in all conditions). 133-AL55 alone is displayed in black; colour codes for each condition are shown in the inset. In the case of CLU (panel C), all pairwise comparisons against 133-AL55 alone were statistically significant (inset) except for the 500:1 CLU curve. In the case of apoE3, all four pairwise comparisons against 133-AL55 alone are significant.

### ApoE3 and CLU alter the fibrillogenesis kinetics of the amyloidogenic LC fragments

To assess the influence of specific exogenous factors on LC aggregation, we performed fibrillogenesis assays in the presence of substoichiometric amounts of amyloid-signature proteins CLU and apoE, at concentrations ranging from 2 µM to 0.02 μM (corresponding to LCs over CLU/apoE3 molar ratios ranging from 5:1 to 500:1). Given the lack of proven association between apoE genotype and AL amyloidosis (28), we tested apoE3, the product of the most common allele. Neither CLU nor apoE3 alone form amyloid fibrils in vitro under the same experimental conditions (data not shown).

Representative fibrillogenesis kinetics are displayed in Figures 1 and 2 (panels A and B) and show that the two chaperones significantly modulate AL aggregation in a dose-dependent manner. At the highest concentration (2 µM, i.e. 5:1 LCs to apoE3/CLU ratio), both apoE3 and CLU completely prevent LC fibrillogenesis over the entire observation time (70 hours). At a tenfold lower dose (0.2 µM, i.e. 50:1 LC to apoE3/CLU molar ratio), protection is still highly significant, but more pronounced for apoE3. In fact, whereas 50:1 CLU delays but does not abolish amyloid formation, 50:1 apoE3 allows fibrillogenesis to occur only in a few reaction wells after a prolonged lag-phase. However, at lower concentrations of chaperones, their effects differ across LCs. Concerning 128-ALH7, the delaying action of apoE3 and CLU is no longer present at ≤250:1 molar ratios (cofactors ≤ 0.04 µM), being replaced by an opposite effect of amyloidogenesis promotion, with significant lag phase shortening (especially pronounced for CLU). In the case of 133-AL55, instead, the protective effect of apoE3 and CLU is still present, albeit reduced, at the 250:1 and - only for apoE3 - at the 500:1 doses, without paradox lag phase shortening. The lag phase variations are summarized in Figures 1 C and D, and 2 C and D, where experimental data are visualized using Kaplan-Meier curves (counting the starting time of fibril exponential growth as the event of interest) (29). Considering all technical and experimental replicates, the ThT amplitude at plateau is not statistically different in the presence of cofactors, except for 128-ALH7 incubated with 50:1 CLU, which displays lower ThT values compared to 128-ALH7 alone.

### CLU and apoE3 are incorporated in the LC fibrillar aggregates

Fibrillogenesis mixtures were centrifuged to recover insoluble aggregates, followed by morphological and biochemical characterization. These tests were performed after 40 h of incubation, to avoid the variability in fibrillogenesis outcomes observed for some conditions at longer times (at this timepoint, in fact, ThT fluorescence is at a plateau in all cases, except for 5:1 CLU/apoE3 and 50:1 apoE3, in which all replicates are always ThT-negative). In all ThT-positive samples, SDS-PAGE analysis and supernatant quantification show practically complete precipitation of LCs, while these are still soluble in the ThT-negative conditions (Figure 3 A-B and Figure S2). Dot blot analysis was then performed to assess the presence of apoE3 and CLU in the insoluble material. Regarding apoE3, this protein is detectable in ThT-negative pellets at the highest tested concentration (2 μM, 5:1 ratio), showing that apoE3 itself can partly aggregate amorphously when present in sufficiently high amounts (Figure 3 C-D). No apoE3 precipitate is instead present in the ThT-negative 50:1 sample, while it is again detectable in the ThT-positive 250:1 and 500:1 pellets, indicating its incorporation in or association with the amyloid aggregates. Regarding CLU, no significant amorphous aggregation of this protein occurs in the ThT-negative specimens. In contrast, it is detectable in the pellets from all ThT-positive conditions, indicating its incorporation in or association with amyloid aggregates. Finally, to verify if apoE3 and CLU can bind to preformed amyloid, we co-incubated these two proteins (2 μM) with naked fibrils from either LC (Figure 3 E-F). The amount of apoE3 and CLU in the insoluble pellets increases in the presence of LC fibrils, showing that both chaperones can also associate to preformed amyloid.

**Figure 3.**
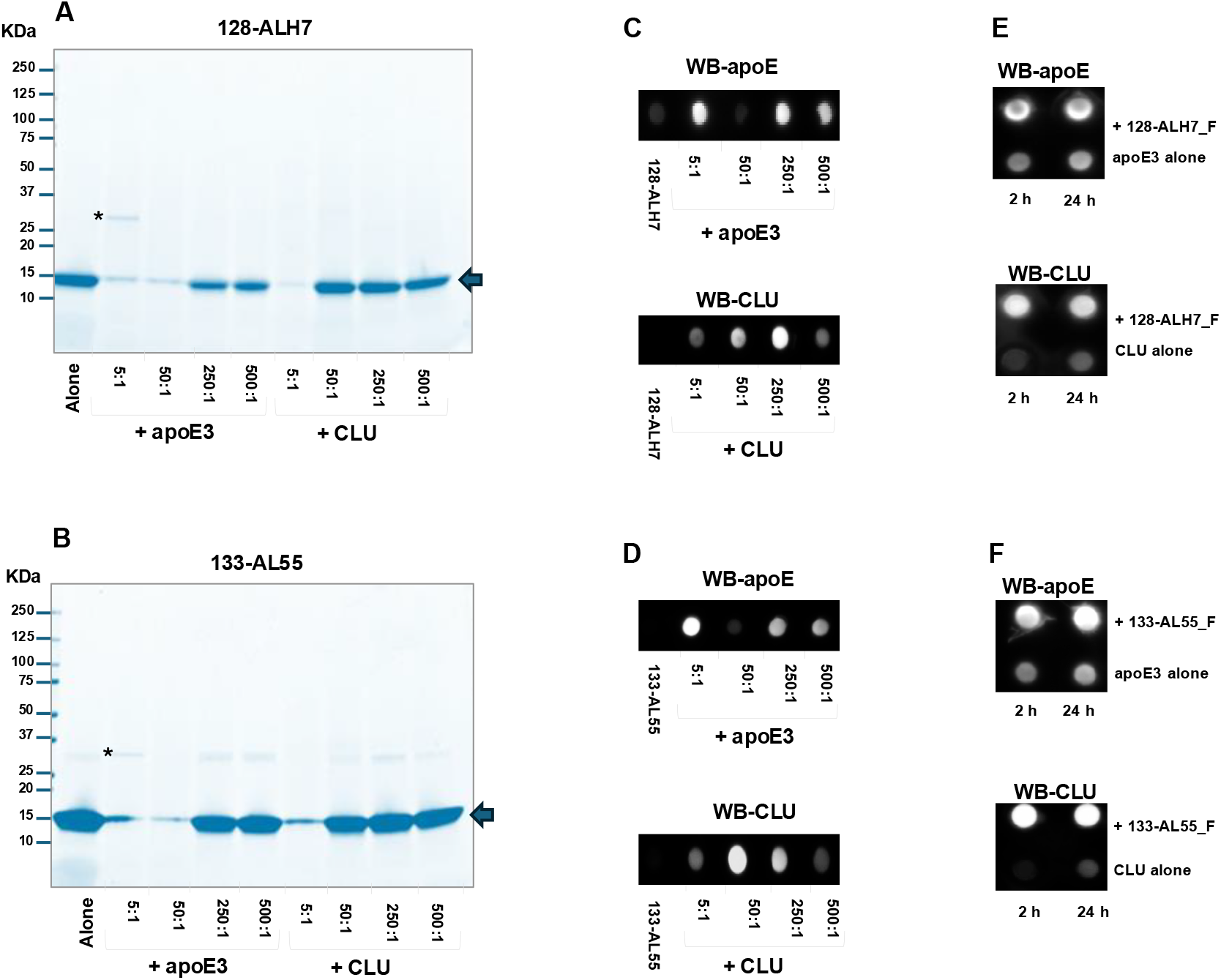
Composition of pellets and supernatants from the LC fibrillogenesis assays in the presence of apoE3 and CLU. (A and B): SDS-PAGE analysis of the aggregates from the incubation of, respectively, 128-ALH7 and 133-AL55 (10 µM) with different substoichiometric concentrations (5:1, 50:1, 250:1 and 500:1 LCs:cofactors molar ratios) of apoE3 or CLU. Each lane contains the material pelleted from 200 μl of suspension collected after 40 h of incubation. Bands corresponding to the LCs are indicated by arrows. The asterisk indicates apoE3. Data are representative of three independent experimental replicates. (C and D): Dot blot detection of CLU and apoE in the pellets from the fibrillogenesis experiments shown in panels A and B, referring respectively to 128-ALH7 and 133-AL55 (data representative of three experimental replicates; material pelleted from 100 μl of suspension); (E and F): Dot blot detection of CLU and apoE bound to preformed amyloid fibrils (indicated as F) of 128-ALH7 and 133-AL55, with which they were incubated for 2h or 24h before aggregate recovery and washing (data are representative of three experimental replicates; material pelleted from 100 μl of suspension).

### Light chain amyloid fibrils grown in the presence of cofactors are morphologically different

Morphological and thermodynamic studies of amyloid aggregates were performed on selected conditions, i.e. those in which ThT increase in the presence of chaperones was reproducibly observed, at the high and low ends of the concentration range. These conditions include 50:1 and 500:1 CLU and 500:1 apoE3. At AFM, 133-AL55 (Fig 4 A-F) exhibit fibrillar aggregates intertwined together (Fig. 4A) with a height of 7.8 ± 0.4 nm. A qualitatively similar morphology is observed in the presence of 500:1 apoE3 (Fig. 4B) and of 50:1 and 500:1 CLU (Fig. 4 C-D), but the corresponding fibril height distributions tend to be shifted to smaller values (Fig. 4 E and F): 6.6 ± 0.4 nm with 500:1 apoE3; 7.4 ± 0.3 nm with 500:1 CLU and 5.5 ± 0.3 nm at 50:1 CLU. The height distribution of fibrils formed in the presence of 50:1 CLU is significantly different (p<0.001) from that of 133-AL55 alone. Regarding 128-ALH7, fibrils have a significantly different morphology compared to 133-AL55, being shorter and tightly packed in massive aggregates (Fig. 4 G), and allowing height measurement only on those protruding from the edge of the clumps. The height of fibrils from 128-ALH7 alone is 3.3 ±0.1 nm. A similar behaviour is found in the presence of cofactors (Fig. 4 H-J); however, with 500:1 apoE, in addition to massive fibril clumps, also isolated globular aggregates and very short fibrils are present (Fig. 4 H). As in 133-AL55, the presence of cofactors affects the fibril height. Yet, in contrast to 133-AL55, incubation with cofactors significantly shifts the height distributions to larger values in all cases (p<0.001) (4.8 ±0.3 nm in the presence of 500:1 apoE3; 6.1 ± 0.4 nm in the presence of 500:1 CLU, 8.5 ± 0.4 nm in the presence of 50:1 CLU) (Fig. 4 K and L).

**Figure 4.**
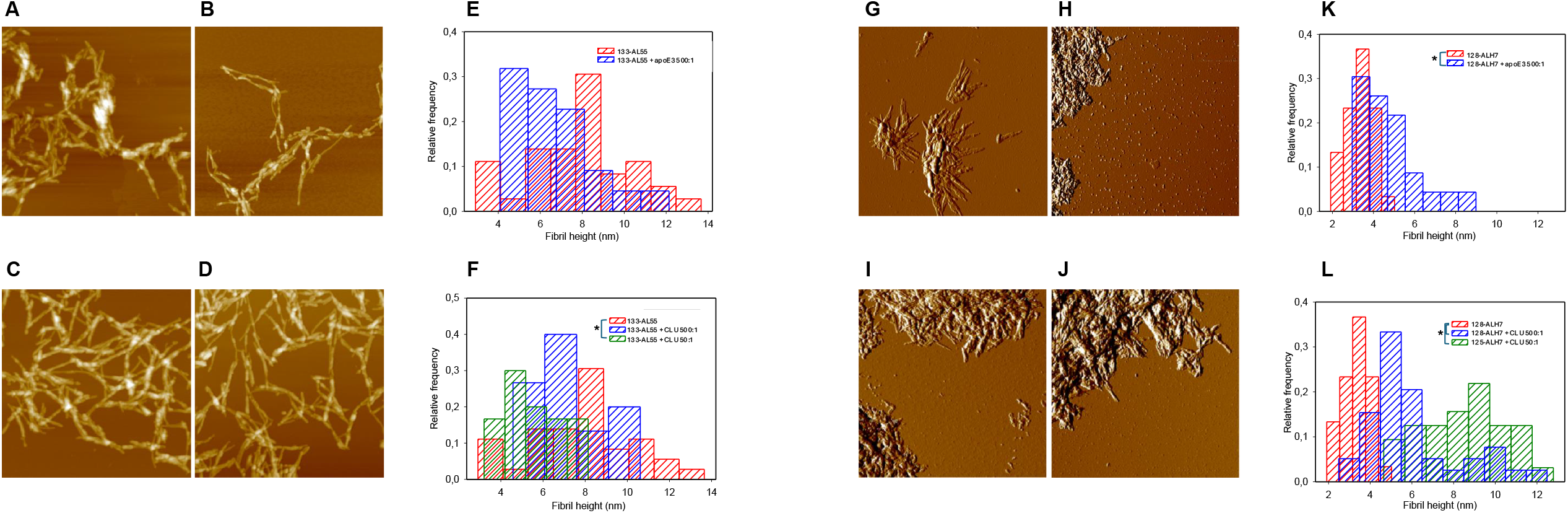
AFM analysis of fibrils formed by 128-ALH7 and 133-AL55 in the presence of apoE3 and CLU. Panels A-D: AFM images (height data) of 133-AL55 aggregated in the absence (A) and in the presence of 0.02 µM (500:1 LCs:cofactors molar ratio) apoE3 (B), 0.02 µM (500:1 ratio) CLU (C), 0.2 µM (50:1 ratio) CLU (D). Scan size 2.5 µm, Z range 70 nm (A, C) and 50 nm (B, D). Panels E and F: height distributions of 133-AL55 fibrillar aggregates in the absence and in the presence of apoE3 (E) and in the absence and presence of CLU (F) at the different concentrations examined. Panels G-J: AFM images (amplitude data) of 128-ALH7 aggregated in the absence (G) and in the presence of 500:1 apoE3 (H), 500:1 CLU (I), 50:1 CLU (J). Scan size 2.5 µm. Panels K and L: height distributions of 128-ALH7 fibrillar aggregates in the absence and in the presence of apoE3 (K) and in the absence and presence of CLU (L) at the different concentrations examined. Significantly different distributions (Mann-Whitney test) are indicated with asterisks.

### Modulation of the thermodynamic stability of LC fibrils grown in the presence of apoE3 and CLU

Changes in thermodynamic stability of amyloid fibrils grown in the absence and presence of apoE3 (500:1 LC to apoe3/CLU molar ratio), and CLU (50:1 and 500:1 molar ratios) were studied in response to increasing concentrations of guanidine isothiocyanate (GdnSCN), after overnight equilibration at 25 °C, 32 °C, 37°C, 41 °C and 45 °C (Fig. 5 A and B). Based on our previous experience with α-synuclein (30), β_2_-microglobulin (31) and TTR fibrils (32), we have applied the same linear polymerization model (Fig. S3 and S4) to derive the free energy of elongation in the absence of denaturant (ΔG_el_^0^), the cooperativity coefficient *m*-value and the midpoint denaturant concentration (C_M_) (Table 1). Depolymerisation curves for 128-ALH7 fibrils show apparent cooperativity in any condition, although this becomes less evident in the presence of 50:1 CLU (Fig. 5B). In contrast, a pre-transition baseline is generally lacking for the unfolding of 133-AL55 fibrils, except in those grown in the presence of 50:1 CLU (Fig. 5A). All fibrils exhibit negative ΔG_el_^0^, both alone and in combination with the interactors. Fibrils from 128-ALH7 alone are more stable than the 133-AL55 counterpart, with significantly more negative ΔG_el_^0^ (p<0.0001) and higher C_M_ values at each temperature (Table 1). The largest reduction in ΔG_el_^0^ is observed for 133-AL55 in the presence of 50:1 CLU and, to a lesser extent, of 500:1 apoE3. On the other hand, only 500:1 apoE3 shows a stabilizing effect on the fibrils formed with 128-ALH7, resulting in a more negative ΔG_el_^0^ compared to all other conditions. As expected, C_M_ values increase accordingly, as stability improves (Table 1).

**Figure 5.**
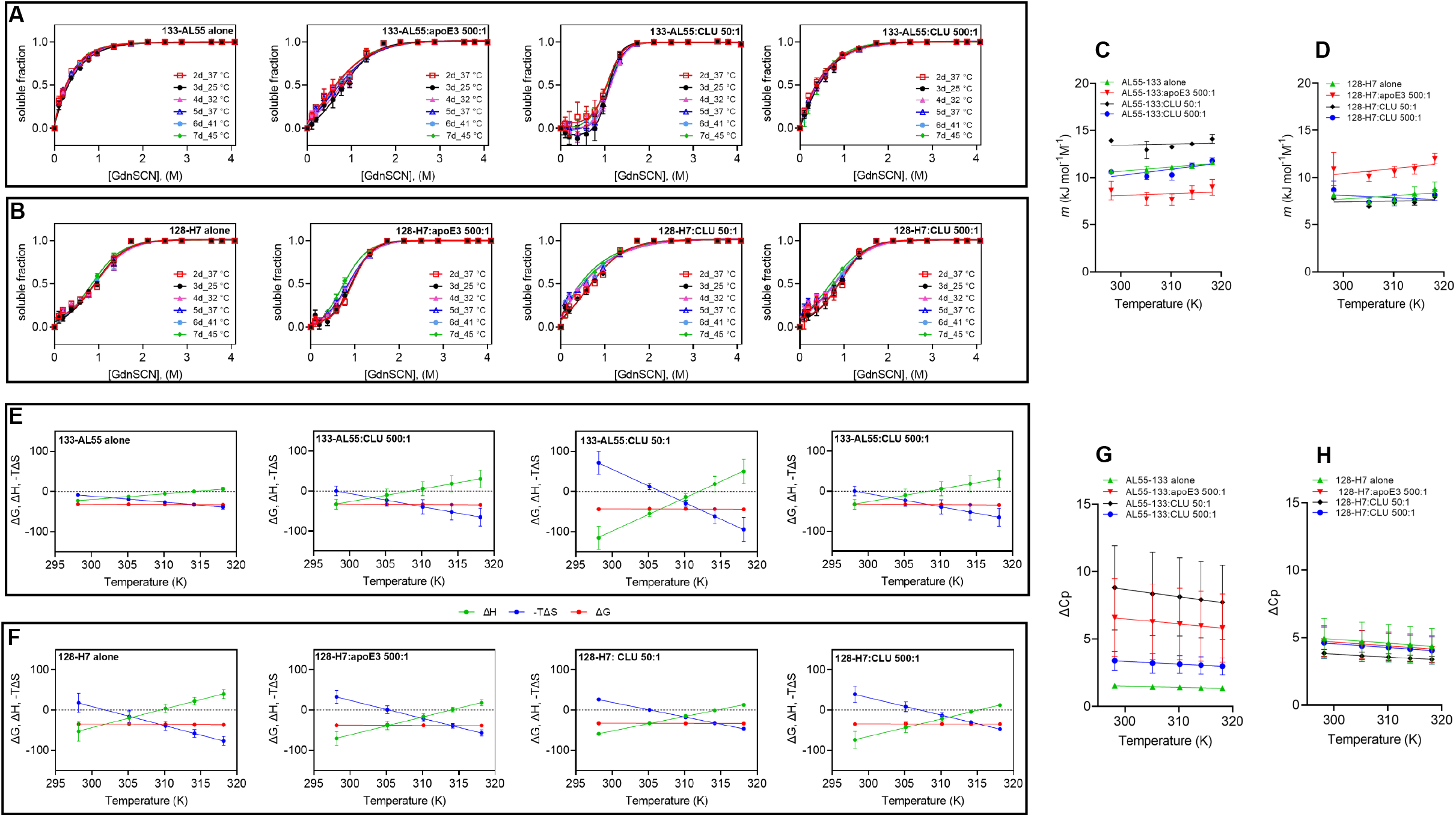
Thermodynamic stability of amyloid fibrils formed by 128-ALH7 and 133-AL55 in the presence of various concentrations of apoE3 and CLU. (A-B) GdnSCN unfolding of 133-AL55 and 128-ALH7 equilibrated at 25 °C, 32 °C, 37 °C, 41 °C and 45 °C. Soluble fraction released during unfolding fitted to a Boltzmann sigmoidal model for demonstrative purpose only. (C-D) Changes in *m* values for 133-AL55 and 128-H7 transitions. (E-F) Changes in ΔH, ΔG_el_^0^ and –TΔS for 133-AL55 and 128-ALH7 fibrils formed alone and in the presence of cofactors. The entropy values were plotted as –TΔS to allow direct comparison with ΔG and ΔH data. (G-H) Changes in heat capacity (ΔCp) for 133-AL55 and 128-ALH7 fibrillar systems in the presence and absence of cofactors.

**TABLE 1.** Thermodynamic stability of 133-AL55 and 128-ALH7 amyloid fibrils.

| $\Delta G_{el}^0$ (KJ mol<sup>-1</sup>) | | | | |
| --- | --- | --- | --- | --- |
| T (K) | 133-AL55 alone | 133-AL55:apoE3 500:1 | 133-AL55:CLU 50:1 | 133-AL55:CLU 500:1 |
| 298.15 | -31.44±0.19 | -34.60±1.12 | -43.79±0.61 | -32.59±0.50 |
| 305.15 | -31.79±0.19 | -34.17±0.81 | -42.62±1.62 | -32.88±0.41 |
| 310.15 | -32.13±0.22 | -34.15±0.55 | -42.90±1.40 | -33.30±0.37 |
| 314.15 | -32.50±0.13 | -34.85±0.59 | -43.75±1.30 | -34.04±0.31 |
| 318.15 | -32.99±0.11 | -35.62±0.79 | -44.35±1.37 | -34.66±0.78 |
| $C_m$ (M) | | | | |
|  | 133-AL55 alone | 133-AL55:apoE3 500:1 | 133-AL55:CLU 50:1 | 133-AL55:CLU 500:1 |
| 298.15 | 0.38±0.01 | 0.84±0.04 | 1.17±0.04 | 0.50±0.05 |
| 305.15 | 0.36±0.01 | 0.81±0.07 | 1.14±0.03 | 0.48±0.04 |
| 310.15 | 0.34±0.01 | 0.75±0.02 | 1.11±0.03 | 0.47±0.02 |
| 314.15 | 0.33±0.01 | 0.72±0.01 | 1.11±0.02 | 0.47±0.02 |
| 318.15 | 0.333±0.003 | 0.72±0.02 | 1.07±0.04 | 0.46±0.05 |
| $m$ (kJ mol<sup>-1</sup> M<sup>-1</sup>) | | | | |
|  | 133-AL55 alone | 133-AL55:apoE3 500:1 | 133-AL55:CLU 50:1 | 133-AL55:CLU 500:1 |
| 298.15 | 10.68±0.24 | 8.63±1.01 | 13.95±0.26 | 10.60±0.27 |
| 305.15 | 10.77±0.34 | 7.73±0.69 | 12.91±0.92 | 10.13±0.32 |
| 310.15 | 11.10±0.14 | 7.65±0.61 | 13.25±0.02 | 10.27±0.51 |
| 314.15 | 11.21±0.02 | 8.41±0.71 | 13.57±0.08 | 11.29±0.30 |
| 318.15 | 11.58±0.23 | 8.97±0.85 | 14.12±0.45 | 11.84±0.31 |
| $\Delta G_{el}^0$ (KJ mol<sup>-1</sup>) | | | | |
| T (K) | 128-H7 alone | 128-H7:apoE3 500:1 | 128-H7:CLU 50:1 | 128-H7:CLU 500:1 |
| 298.15 | -34.96±0.59 | -37.85±1.11 | -32.99±0.25 | -34.90±1.49 |
| 305.15 | -34.67±0.55 | -36.86±0.71 | -32.15±0.20 | -33.83±0.81 |
| 310.15 | -35.31±0.32 | -37.75±1.29 | -33.04±0.80 | -34.56±1.35 |
| 314.15 | -36.29±0.42 | -38.09±0.71 | -33.05±0.21 | -34.62±0.89 |
| 318.15 | -36.61±0.40 | -38.30±0.69 | -33.43±0.17 | -34.93±0.69 |
| $C_m$ (M) | | | | |
|  | 128-H7 alone | 128-H7:apoE3 500:1 | 128-H7:CLU 50:1 | 128-H7:CLU 500:1 |
| 298.15 | 0.91±0.04 | 0.97±0.06 | 0.73±0.00 | 0.88±0.07 |
| 305.15 | 0.91±0.04 | 0.88±0.04 | 0.60±0.01 | 0.79±0.07 |
| 310.15 | 0.90±0.03 | 0.88±0.03 | 0.63±0.01 | 0.80±0.07 |
| 314.15 | 0.91±0.04 | 0.85±0.05 | 0.58±0.01 | 0.76±0.06 |
| 318.15 | 0.85±0.03 | 0.76±0.04 | 0.53±0.02 | 0.70±0.05 |
| $m$ (kJ mol<sup>-1</sup> M<sup>-1</sup>) | | | | |
|  | 128-H7 alone | 128-H7:apoE3 500:1 | 128-H7:CLU 50:1 | 128-H7:CLU 500:1 |
| 298.15 | 8.17±0.32 | 10.90±1.78 | 7.81±0.25 | 8.69±0.97 |
| 305.15 | 7.41±0.41 | 10.12±0.54 | 6.94±0.17 | 7.37±0.51 |
| 310.15 | 7.63±0.48 | 10.59±0.55 | 7.31±0.16 | 7.60±0.64 |
| 314.15 | 8.27±0.73 | 10.85±0.48 | 7.33±0.28 | 7.66±0.62 |
| 318.15 | 8.77±0.77 | 12.03±0.56 | 7.98±0.22 | 8.16±0.57 |

Values of $\Delta G_{el}^0$ (kJ mol<sup>-1</sup>), free energy of elongation in the absence of denaturant, $C_M$ (M), midpoint denaturation and $m$ , cooperativity coefficient were calculated with the linear polymerization mode as described under “Experimental procedures.”

The cooperativity coefficient *m*-values were also considered, as they correlate with changes in accessible surface area upon unfolding (ΔASA) (33). Interestingly, unfolding of 133-AL55 fibrils formed in the presence of 50:1 CLU exhibits higher *m*-values; this is consistent with a more compact fibrillar architecture that exposes a larger surface area upon denaturation and a more cooperative transition compared with unfolding of fibrils by 133-AL55 alone. In contrast, the presence of apoE3 in 133-AL55 fibrillar samples appears to reduce the *m*-value, suggesting a change in the ΔASA associated with a less cooperative unfolding transition. Whilst the *m*-values for 133-AL55 do not change with temperature, the higher cooperativity coefficients values for the unfolding of 128-ALH7 fibrils in the presence of 500:1 apoE3 slightly increase with temperature (Fig. 5 C and D). To quantify the changes in enthalpy (ΔH), entropy (ΔS) and heat capacity (ΔCp) for each condition, ΔG_el_^0^ were converted into the corresponding equilibrium constant (K) to build the Van’t Hoff plot, in which values of lnK vs 1/T were interpolated using a quadratic polynomial equation (34). The temperature dependence of the thermodynamic parameters determined for each fibrillar system in every condition shows clear compensation between enthalpic and entropic contributions, which maintain DΔG near zero across the entire interval (Fig. 5 E and F). ΔH and ΔS changes are consistent with a positive heat capacity change (ΔCp > 0) (Fig. 5 G and H), in which the dominant thermodynamic contribution shifts from enthalpic at low temperatures to entropic at higher temperatures. Whereas ΔH and ΔS changes for 133-AL55 amyloid fibrils are limited across the 25 °C-45 °C range, the temperature dependence of the two thermodynamic components increases for fibrils formed in combination with apoE3 and, particularly, with 50:1 clusterin. The latter system shows the largest ΔCp, concomitantly with a higher *m*-value compared to the fibrils formed by the light chain fragment alone, with both parameters pointing to an expanded ΔASA and a reorganization of the hydration shell. On the other hand, comparison of fibrils formed by 128-ALH7 alone with those assembled in the presence of cofactors reveals only modest variations in thermodynamic parameters.

## DISCUSSION

We have investigated the influence of extrinsic factors in the aggregation of immunoglobulin LCs with intrinsic differences in primary sequence. The current data confirm the previous evidence that LC fragments encompassing the V_L_ and the N-terminal part of the C_L_ (V_L_-C_L_ fragments) convert into amyloid *in vitro* under physiological conditions (5). The fibrillogenesis kinetics, however, depend on the V_L_ sequence: of the two LCs studied herein, 133-AL55 (IGLV6-57) had a significantly shorter lag phase compared to 128-ALH7 (IGLV1-51). The faster aggregation of 133-AL55 is in line with the computational predictions of more APRs in this LC, two of which (residues 17-21 and 31-36), located in the V_L_, are agreed upon by all the tested algorithms. While the former APR is engaged in the β-strand within a conserved V_L_ β-sheet, residues 31-36 belong to an exposed loop that may be especially prone to establishing intermolecular interactions upon conformational fluctuations.

Our data, however, show that fibrillogenesis of V_L_-C_L_ LC fragments is also significantly modulated by heterotypic factors. Specifically, we have explored apoE and CLU, two amyloid signature proteins serving as diagnostic markers of AL and of virtually all other systemic amyloidoses (35). The structure and biochemical properties of apoE and CLU tie up with their ability to interact with misfolded proteins. Clusterin (apoJ) is a glycosylated 60 kDa disulfide-linked heterodimer, secreted by many cell types and found in all extracellular fluids, with a plasma concentration of 30-135 μg/mL (36). Its structure has a three-domain architecture with two disordered hydrophobic peptide tails (37, 38). This protein is implicated in various processes, including complement regulation, apoptosis and lipid transport. Clusterin is one of the most important extracellular holdases, able to bind misfolded proteins through interaction with exposed hydrophobic regions and formation of high-molecular weight complexes, thus inhibiting aggregation and directing nonnative clients to receptor-mediated cellular degradation (12, 13, 23, 29). ApoE is a water-soluble 34-kDa apolipoprotein involved in lipid transport in plasma and central nervous system, predominantly produced by liver and astrocytes. In humans, three major apoE alleles exist (ε2, ε3 and ε4), ε3 being the most common. ApoE circulates in plasma on high-density and very low-density lipoproteins with a concentration of 60–120 μg/mL and mediates the cellular uptake and trafficking of these particles. ApoE exists in equilibrium between a lipid-bound and a lipid-free form and has a structure organized in multiple amphipathic α-helices that can interact with lipids and hydrophobic particles through exposed hydrophobic patches (19).

The pathophysiological role of extracellular chaperones in systemic amyloidoses has been scarcely explored, while more evidence is available for cerebral forms. In Alzheimer’s disease, the importance of scrutinizing apoE and CLU is strongly supported by the clinical evidence that gene variants of these two proteins are major risk factors for late-onset disease (14, 15, 39). *In vitro* and *in vivo* studies show that apoE and CLU bind both to prefibrillar conformers during the nucleation stage and to preformed fibrils, and that substoichiometric concentrations of these interactors modulate Aβ amyloidogenesis (19, 37, 40, 41). Molecular simulations suggest apoE binding to hydrophobic surfaces on sides or ends of Aβ fibrils, interfering respectively with fragmentation and secondary nucleation, or with elongation and depolymerization. The effects of CLU and apoE on Aβ amyloidogenesis, however, are concentration dependent but not necessarily mono-phasic. At sufficiently high ratios of CLU/apoE to Aβ peptide, in fact, both interactors exert a significant inhibitory effect (29, 41, 42), while amyloidogenesis can be favoured by low chaperone concentrations, possibly due to stabilization of transient on-pathway species (13, 22). No such epidemiological association with CLU/apoE gene variants, on the other hand, has been demonstrated for systemic amyloidoses. The apoE genotype was ruled out as a risk factor in AL or in ATTR, and no evidence is available on CLU gene variants in these diseases (28). A functional role of CLU was nevertheless indicated by pivotal studies showing decreased CLU serum levels in AL and ATTR amyloid cardiomyopathy patients (43), and significant alterations in the glycosylation profile of this protein in ATTRwt, suggesting reduced chaperoning efficiency (43-48).

Herein, we demonstrate that both apoE3 and CLU significantly affect the kinetics and outcome of LC fibrillogenesis, with complex effects that range from anti-to pro-amyloidogenic depending on their concentration and on the LC’s intrinsic amyloidogenic propensity. In the case of fast-fibrillating 133-AL55 LC, both CLU and apoE3 exert exclusively a protective effect, manifesting as a dose-dependent lag phase increase, while in 128-ALH7 this delay occurs only when the interactors’ concentration is sufficiently high. In tissues, where cells participate in proteostasis by removing chaperone-client complexes, this delay provides the opportunity to clear harmful precursors before amyloid accumulation. However, in slow-fibrillating 128-ALH7, a paradox effect of fibrillogenesis anticipation was exerted by low amounts of apoE3 and, even more markedly, of CLU. These differences between LCs subtend precursor-specific equilibria between folded and misfolded monomers, oligomeric aggregates and amyloid. For those LCs in which low cofactors concentration accelerates amyloid aggregation, conditions that reduce interstitial chaperones availability *in vivo* (*e*.*g*. low serum levels or sequestration by other partners) may in fact further amplify the amyloidogenesis risk.

The thermodynamic analysis of amyloid fibrils demonstrates that extracellular chaperone proteins modulate both fibril stability and the assembly mechanism. Although all systems show negative ΔG values, indicating spontaneous elongation, differences are observed in both the magnitude of ΔG and depolymerisation profiles. Notably, 128-ALH7 fibrils exhibit cooperative behaviour consistent with a nucleation–elongation mechanism, while 133-AL55 fibrils exhibit cooperativity in the presence of interactors, particularly CLU. Deviations from ideal sigmoidal transitions suggest intermediate species and more complex equilibria, supporting a model in which heterotypic interactions reshape the aggregation pathway. Modulation of ΔG by apoE3 and CLU is strongly dependent on sequence and chaperone concentration. CLU at 0.2 μM induces substantial stabilisation of 133-AL55 fibrils, whereas apoE3 selectively stabilises 128-ALH7 at low concentration. These effects are accompanied by changes in the cooperativity coefficient *m*, which reflects the variation in solvent-accessible surface area (ΔASA) upon unfolding. Notably, the increase in m values observed for 133-AL55 in the presence of CLU 0.2 μM indicates a more cooperative transition and suggests the formation of structurally more compact or better-packed fibrillar assemblies. In contrast, the decrease in *m* induced by apoE3 indicates distinct structural rearrangements involving reduced cooperativity. Importantly, changes in *m* are not always accompanied by proportional variations in ΔCp, indicating that alterations in exposed surface area and solvent interactions are partially decoupled. This suggests that cofactor-induced structural rearrangements involve not only differences in fibril packing but also changes in hydration and solvent organisation at the fibril interface. In this context, the positive ΔCp values observed under all conditions indicate a significant contribution of solvent reorganisation to fibril stability, a behaviour that, although uncommon, has been reported for specific amyloid systems such as α-synuclein (49) and insulin fibrils (50). Mechanistically, the concentration-dependent effects of CLU and apoE align with their preferential interaction with prefibrillar intermediates rather than monomeric species. At high concentrations, CLU and apoE likely sequester aggregation-prone intermediates into soluble high-molecular-weight complexes, preventing nucleation and fibril growth. At lower, substoichiometric concentrations, partial sequestration slows aggregation while allowing fibril formation to proceed via alternative pathways, ultimately favouring the selection of more stable polymorphic states. This interpretation is supported by the increased lag phase observed under these conditions and by the thermodynamic signatures of the resulting fibrils.

Analysis of the aggregates shows that both CLU and apoE3 are co-deposited with amyloid fibrils and our experiments show that these two proteins can interact with LCs at distinct timepoints, i.e. with prefibrillar species, affecting amyloidogenesis kinetics, and with preformed fibrils. Notably, however, the incorporation of CLU and apoE3 into aggregates even at very low concentrations suggests that the interactions occurring in the early aggregation stages may not be limited to transient kinetics modulation but could contribute structurally to the final assemblies. This hypothesis finds support in the AFM evidence that the fibril height is affected by apoE3 and CLU. In all ThT-positive samples, LCs aggregate virtually completely, with only amyloid fibrils visible by AFM except in 128-ALH7 incubated with 0.02 μM (500:1 molar ratio) apoE3, in which also small amounts of globular structures are present. The morphology of the two LCs fibrils is, however, significantly different: aggregates from 133-AL55 are longer, while 128-ALH7 ones are short and needle-like, suggesting a higher rate of fragmentation and secondary nucleation. Amyloids grown in the presence of cofactors maintain such qualitative features but vary significantly in height distributions compared to their untreated counterparts. The heights measured by AFM are consistent with those expected for single or multiple filaments fibrils (51), suggesting that the two LCs can form polymorphic fibril populations composed of a different number of protofibrils, whose relative abundance is shifted by the heterotypic interactors. In particular, while the cofactors shift the height distributions of 133-AL55 fibrils to lower values, the opposite occurs in 128-ALH7, indicating once more that the result of the heterotypic interactions varies according to the individual properties of the amyloidogenic precursor.

These findings support a model in which extracellular chaperones act as modulators of the amyloid energy landscape, influencing both the kinetics and thermodynamic stability of fibril formation. Rather than simply stabilising or destabilising fibrils, apoE3 and CLU promote the selection of distinct polymorphic states characterised by different packing, solvent exposure, and cooperativity. This dual role, combining kinetic control and structural selection, may contribute to the heterogeneity (52, 53) and persistence of amyloid deposits *in vivo*.

Translationally, our results support a Janus-like role of chaperones in AL amyloidosis, which can turn from protective to noxious depending on their levels in sites where misfolding and aggregation take place. Levels of CLU and lipid-free apoE in the extracellular space are expectedly lower than those measurable in plasma, and oligomerization, as in the case of CLU, may further reduce interstitial transfer (54). Fluctuations in serum concentration or decline with aging could thus pose a realistic risk for not achieving protective levels. We also show that the interactors’ effects are not identical on different LCs, likely contributing to the phenotypic variability of this disease. Tuning the extracellular chaperone system could be critical to modulate AL amyloidosis and efforts are needed to further characterize the effects of heterotypic LC interactors on a population-based scale.

## EXPERIMENTAL PROCEDURES

### Chemicals and reagents

All chemicals were from Sigma-Aldrich Merck (Darmstadt, Germany), Thermo Fisher Scientific (Pittsburgh, PA, USA) or VWR International (Radnor, PA, USA), unless otherwise specified. Recombinant human apoE3 was purchased from Peprotech/Gibco (catalog # 350-02, Thermo Fisher Scientific).

### Protein expression and purification

Light chain fragments extending from the N-terminus to position 22 of the C_L_ were produced from the sequences of amyloidogenic human immunoglobulin λ light chains (LCs) AL-55 (IGLV6-57 germline, GenBank #MH670901) and AL-H7 (IGLV1-51 germline, GenBank #KC433671). These two fragments, which are composed of, respectively, 134 and 129 amino acids (including the N-terminal methionine), are indicated as 133-AL55 and 128-ALH7. The LC sequences, cloned in pET-29a vectors between the NdeI and XhoI restriction sites, were expressed in BL21(DE3) E. coli as inclusion bodies, solubilized as previously described (5). The purification protocol used for 133-AL55 was reported in detail and includes refolding in the presence of reduced and oxidized glutathione (GSH), followed by ion exchange and size-exclusion chromatography (5). In the case of 128-ALH7, a first size-exclusion chromatography step was performed using a Superdex HiLoad 26/600 75pg column (Cytiva, Sigma-Aldrich) in 20 mM NaHPO_3_, pH 7.4, 5 M GdnHCl. The eluted LC fractions were refolded by dropwise by addition of the same refolding buffer used for 133-AL55 (0.1 M NaHPO_3_ pH 7.4, 1 mM EDTA, 1 mM PMSF, 5 mM reduced L-glutathione, and 0.5 mM oxidized L-glutathione), followed by dialysis against progressively lower NaCl concentrations (dialysis A: 20 mM NaHPO_3_ / 150 mM NaCl; dialysis B: 20 mM NaHPO_3_ / 75 mM NaCl; dialysis C: 20 mM NaHPO_3_ / 37,5 mM NaCl; dialysis D: 50 mM Tris-HCl). The resulting protein was further purified by ion exchange chromatography on an HiPrep Q HP 16/10 (Cytiva, Sigma-Aldrich) and dialyzed against PBS. The purified LCs were characterized by reducing and non-reducing SDS-PAGE, mass spectrometry and circular dichroism. Recombinant clusterin containing a small EPEA tag at the C terminus of the β chain was produced in HEK293-F cells following the protocol described in (55), with minor adaptations (56). Briefly, suspension growing HEK293-F cells (Life Technologies, UK) were transfected at of 10^6^ cells/ml^-1^, using a mixture of 0.8 µg of plasmid DNA containing the clusterin gene (rCLU-CT_pcDNA3.1(+)), 0.2 µg of a pUPE expression plasmid for mammalian expression (U-Protein Express BV) carrying the human cDNA of the pro-protein convertase furin, and 5 µg of linear polyethyleneimine (PEI; Polysciences). Cells were harvested 6 days after transfection by centrifuging the medium for 15 minutes at 1000 x g. The clarified conditioned medium was subject to a high-speed centrifugation 15 minutes at 4000 x g, followed by flash-freezing in liquid nitrogen and storage at -80 °C until usage. Recombinant CLU was purified using a 1 ml C-tag affinity column (Capture Select, Thermo Fisher Scientific), eluted with 2 M MgCl_2_ in 20 mM Tris-HCl buffer, pH 7.4, dialyzed against PBS and characterized by SDS-PAGE. Protein concentrations were determined spectrophotometrically at 280 nm using the respective predicted molar extinction coefficients. Purified proteins were stored frozen at -80 °C.

### Fibrillogenesis experiments

All *in vitro* fibrillogenesis and co-fibrillogenesis studies were performed under physiological conditions of pH (7.4), temperature (37 °C) and ionic strength (PBS), with agitation (900 rpm). Amyloid aggregation was monitored over time (15-min intervals measurements) following the increase in Thioflavin T fluorescence (ThT; working concentration 10 μM) (excitation wavelength 445 nm; emission 480 nm), in sealed 96-well black-walled plates (Corning, NY, USA), on a Clariostar microplate reader (BMG Labtech, Ortenberg, Germany) (5). LC fragments were used at a concentration of 10 µM, whereas co-fibrillogenesis studies were performed with different concentrations of apoE3 or CLU (2 μM, 0.2 μM, 0.04 μM and 0.02 μM, corresponding to LCs to cofactors molar ratios of 5:1, 50:1, 250:1 and 500:1). At least three full experimental replicates for all assays were performed; each experimental replicate included 3 to 5 technical replicates (i.e. microplate wells) per condition. Kinetics were visualized using GraphPad Prism. Insoluble aggregates to be used for biochemical and morphological characterization were separated from soluble proteins by centrifugation (14,000 rpm, 20 min, room temperature), followed by washing with 1 ml of PBS. Binding of CLU and apoE3 to preformed LC fibrils was studied by adding each protein (2 μM) to 100 µl of 10 µM fibrils from 128-ALH7 or 133-AL55, followed by incubation at 37 °C under agitation for 2h or 24h. Insoluble aggregates were recovered as described above and washed once with PBS prior to western blot analysis.

### Sodium dodecyl sulphate polyacrylamide gel electrophoresis (SDS-PAGE) and dot blot

Electrophoresis and western blotting were performed under denaturing and reducing conditions as previously described, on 4-20% polyacrylamide gradient mini gels (Bio-Rad, Hercules, CA, USA) (5), followed by staining with colloidal Coomassie blue G-250 (GelCode Blue Stain Reagent, Thermo Fisher Scientific) and imaging using a Bio-Rad ChemiDoc MP system. Gel bands intensity was measured on digitized images using ImageLab software (Bio-Rad). For dot blot analysis, protein pellets were resuspended in 8 M urea to dissolve the aggregates, transferred onto PVDF membranes and probed with mouse monoclonal anti-clusterin α-chain, clone 41D (Sigma-Aldrich Merck) or mouse monoclonal anti-apoE (Santa-Cruz Biotechnology, Dallas, Texas, USA) primary antibodies.

### Atomic Force Microscopy (AFM)

133-AL55 and 128-ALH7 samples (10 μM) aggregated in the absence and in the presence of different concentrations (0.2 and 0.02 μM, i.e. 50:1 and 500:1 LC:cofactors molar ratios) of CLU and apoE3 were centrifuged at 4000 rpm for 5 min and pellets were collected. Each pellet aliquot was resuspended in ultrapure water at its original concentration. For AFM inspection, 10 μL of the aggregate suspension were deposited onto a freshly cleaved mica substrate (1.0 cm x 1.0 cm) and dried overnight under vacuum at room temperature. AFM images were acquired in tapping mode in air, using a Dimension 3100 SPM (Bruker, Karlsruhe, Germany) equipped with a “G” scan head (maximum scan size 100 μm) and driven by a Nanoscope IIIa controller (Bruker). Rectangular Al-coated cantilevers (TESPA-V2, Bruker) with nominal spring constant of 40 N/m, nominal resonance frequency of 300 kHz and nominal tip radius of 7 nm were employed. The amplitude setpoint was set to 80% of the free oscillation amplitude, and the scan rate was 0.4-1.2 Hz. Fibril heights were measured from the cross sections of topographic AFM images.

### Thermodynamic characterisation of Amyloid Fibrils

The thermodynamic stability of LC aggregates formed in absence or presence of cofactors was quantified using a microplate assay, based on ThT fluorescence measurement in the presence of increasing concentrations of denaturant (guanidine isothiocyanate, GdnSCN) (57). Briefly, stock aggregates of 128-ALH7 and 133-AL55 were prepared in PBS under the following conditions, selected based on the kinetics studies because consistently characterized by significant lag-phase modifications: (1) 30 µM LC alone; (2) 30 µM LC + 0.6 µM CLU (50:1 molar ratio); (3) 30 µM LC + 0.06 µM CLU (500:1 ratio); (4) 30 µM LC + 0.06 µM apoE3 (500:1 ratio). Incubation was carried out for 72 hours in all cases. Aliquots of stock solutions were withdrawn for ThT measurement and characterization of aggregates (isolated by centrifugation, washed as described above and solubilized in 8 M urea for 1h) by SDS-PAGE and protein quantification by Bradford assay. The aggregate stability assay was performed in Costar 96-well black-walled plates; briefly, aggregates suspensions were diluted to a concentration of 9.6 µM in wells containing increasing concentrations of GdnSCN (ranging from 0 to 4.1 M) and 10 μM ThT in PBS. Plates were sealed and sequentially equilibrated overnight at different temperatures (37→25→32→37→41→45 °C). After each overnight incubation, ThT fluorescence was measured in triplicates before moving to the next temperature. Three replicates of each dilution series were performed for each condition.

### Determination of thermodynamic parameters

ThT fluorescence values at each denaturant concentration were normalized to the value of the corresponding sample in the absence of denaturant [M_fib_]. The depolymerized fraction derived as:

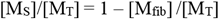

with [M_T_], total amount of protein and [M_S_],soluble fraction at each denaturant concentration was plotted with GdnSCN concentrations and analyzed using the linear polymerization model (32, 58):

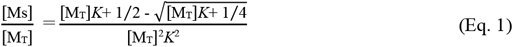

In the presence of chemical denaturants, i.e. GdnSCN, the equilibrium constant *K* can also be expressed as:

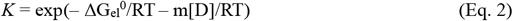

in which ΔG_el_^0^ is the free energy of elongation in the absence of denaturant, *m* is a cooperativity coefficient, R is the gas constant and T is the absolute temperature. By introducing these terms in the equation 1, values of ΔG_el_^0^ and *m* values were determined at each temperature using KaleidaGraph 4.0 (Synergy Software, Reading, PA) together with the midpoint denaturation for each condition. ΔG_el_^0^ values were then converted to the corresponding ln*K* to construct the Van’t Hoff plots (ln*K* vs 1/T). The resulting graphs were then analysed with GraphPad Prism 9.5.1 according to the quadratic polynomial equation:

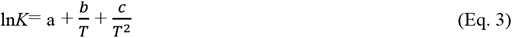

and enthalpy 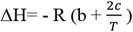, entropy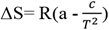 and heat capacity 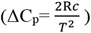 changes derived for each temperature (Haidacher D PNAS 1996). All measurements are reported as mean ± SD for each of the 3 series of samples.

### APR analysis

Aggregation-prone regions in the sequences of 133-AL55 and 128-ALH7 were predicted using multiple models: AmylPred2 (with consensus of at least 4 distinct algorithms), ANuPP and Aggregscan. A score ranging from 0 to 3 was calculated for each amino acid residue based on the number of models predicting that position as belonging to an APR. Structures of LC fragments were predicted using AlphaFold3; APRs were mapped on the structures using ChimeraX (v1.10.1) (59).

### Analysis of aggregation lag phase with Kaplan-Meier curves

The kinetics of protein aggregation were analysed using a survival analysis approach (29). Each experimental replicate was treated as an independent observation and the event was defined as the onset of aggregation, identified as the ThT fluorescence increase starting point. For samples that did not exhibit aggregation within the experimental time window, the final time point of the experiment was defined as the event. Curves were estimated using the Kaplan-Meier method and represent the cumulative probability that a sample has not yet aggregated at time t.

### Statistics

The number of experimental and technical replicates is detailed in each experimental procedures section and in figure legends. Differences in mean values were determined by 2-tailed t-test for independent samples and considered statistically significant for p values < 0.05. Differences in aggregation kinetics between experimental conditions (Kaplan-Meier analyses) were evaluated using the log-rank (Mantel-Cox) test with Bonferroni correction. Differences in the height distribution of amyloid fibrils studied by AFM were evaluated by Mann-Whitney test. All analyses were performed using GraphPad Prism 10.6.0.

## Acknowledgements

We are deeply grateful to Prof. Mark Wilson (University of Wollongong) for providing plasmid and protocols for clusterin production. This research was supported by: Italian Ministry for University and Research grants FIS 2021 FIS00001548; Dipartimenti di Eccellenza 2018–2022 and 2023–2027; PRIN 2022 20225HNCZK, PRIN PNRR 2022 P20224WAME); Cariplo grants 2022-0578 and 2022-0535; PNRR grants (PNC “INNOVA” PNC-E3-2022-23683266; CN00000041; PE00000007, INF-ACT; PE0000006, MNESYS); Pfizer (2022-73552599); Telethon Multiround 21-24 (GMR22T1067); AIRC IG Grant 32240; the Ehlers-Danlos Society (Rarer Types Disease Grant 2023), the Giovanni Armenise-Harvard Foundation (CDA 2013); the Italian Ministry of Health (Piano Operativo Salute, IMMUNO-HUB), the Molecular Scale Biophysics Research Infrastructure (MOSBRI, 101004806—H2020-INFRAIA-2018-2020/ H2020-INFRAIA-2020-1). V. B. is Emeritus Professor of Medical Biochemistry at University College London, London (UK). None of the funding sources had roles in study design, collection, analysis, and interpretation of data, in the writing of the report and in the decision to submit this article for publication.

## SUPPLEMENTAL FIGURE LEGENDS

**Figure S1**. (A) Sequences of 128-ALH7 and 133-AL55, aligned using ClustalW. (B) Aggregation-prone regions (highlighted amino acids) predicted using AmylPred2 (consensus of ≥4 distinct algorithms), ANuPP and Aggregscan. The colour indicates the number of models predicting that position as belonging to an APR. (C and D) The APRs shown in B are mapped on the predicted structures of, respectively, 128-ALH7 and 133-AL55.

**Figure S2**. Residual 128-ALH7 (A) and 133-AL55 (B) in the supernatants (10 μl) of the same samples shown in panels A and B of Figure 3 (reducing conditions). Arrows indicate LC bands; asterisk indicates apoE3; § indicates CLU (α/β monomers band and heterodimer). ApoE and CLU are visible only when used at 2 μM concentration. Images are representative of three experimental replicates.

**Figure S3**. Soluble fraction released from 133-AL55 fibrils at increasing GdnSCN concentrations was analysed with Equation 1 following the linear polymerization model as described under Experimental Procedures. Curves shown for each sample (A, B, C) as mean and SD of three readings.

**Figure S4**. Soluble fraction released from 128-ALH7 fibrils at increasing GdnSCN concentrations was analysed with Equation 1 following the linear polymerization model as described under Experimental Procedures. Curves shown for each sample (A, B, C) as mean and SD of three readings.

