## Supplemental figures for "Extracellular chaperones modulate AL light chains fibrillar aggregation and contribute to amyloid structural heterogeneity"

**Figure S1**

**A**

```
128-ALH7   QSVLTQPPSVSAAPGQKVTISCS----NVGKNFVSWYQQFPGTAPKVVIYDTDKRPSDIP
133-AL55   NFMLTQPHSVSESPGKTLTISCTGSSASIASHYVQWYQQRPGGAPTTLIYENDQRPSEVP
           : :**** *** :**:.:****:    .:..::*.**** ** **..:**:.*:***::*

128-ALH7   DRFSGSK--SGTSATLDITGLQTGDEADYYCGTWDSGLNGGVFGGGTKVTVLGQPKAAPS
133-AL55   DRFSGSIDSSSNSASLTISGLKTEDEADYYCQSYD-GNNHWVFGGGTKLTVLSQPKAAPS
           ******   *..**:* *:**:* ******* ::* * *  *******:***.*******

128-ALH7   VTLFPPSSEELQAN
133-AL55   VTLFPPSSEELQAN
           **************
```

**B**

128-ALH7

QSVLTQPPSVSAAPGQKVTISCSNVGKNFVSWYQQFPGTAPKVVIYDTD
KRPSDIPDRFSGSKSGTSATLDITGLQTGDEADYYCGTWDSGLNGGVFG
GGTKVTVLGQPKAAPSVTLFPPSSEELQAN

133-AL55

NFMLTQPHSVSESPGKTLTISCTGSSASIASHYVQWYQQRPGGAPTTLI
YENDQRPSEVPDRFSGSIDSSSNSASLTISGLKTEDEADYYCQSYDGNN
HWVFGGGTKLTVLSQPKAAPSVTLFPPSSEELQAN

| |
|---|
| 0 |
| 1 |
| 2 |
| 3 |

**C**

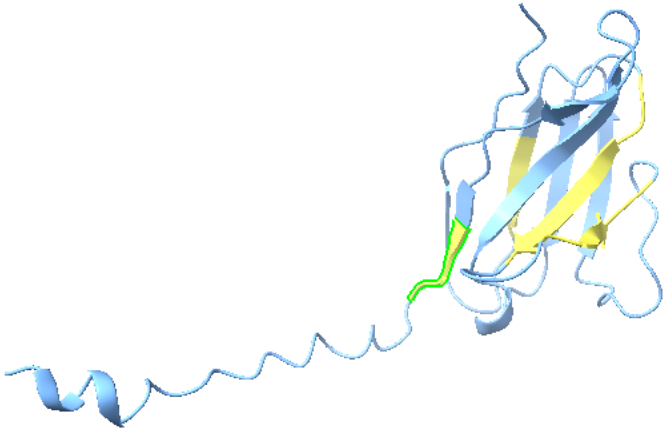

**128-ALH7**

**D**

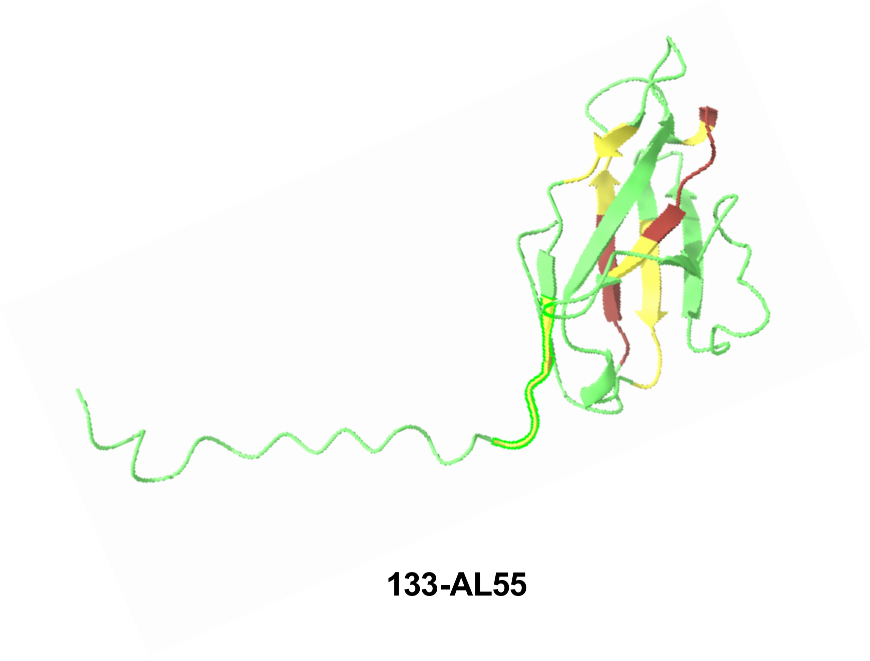

**133-AL55**

Figure S2

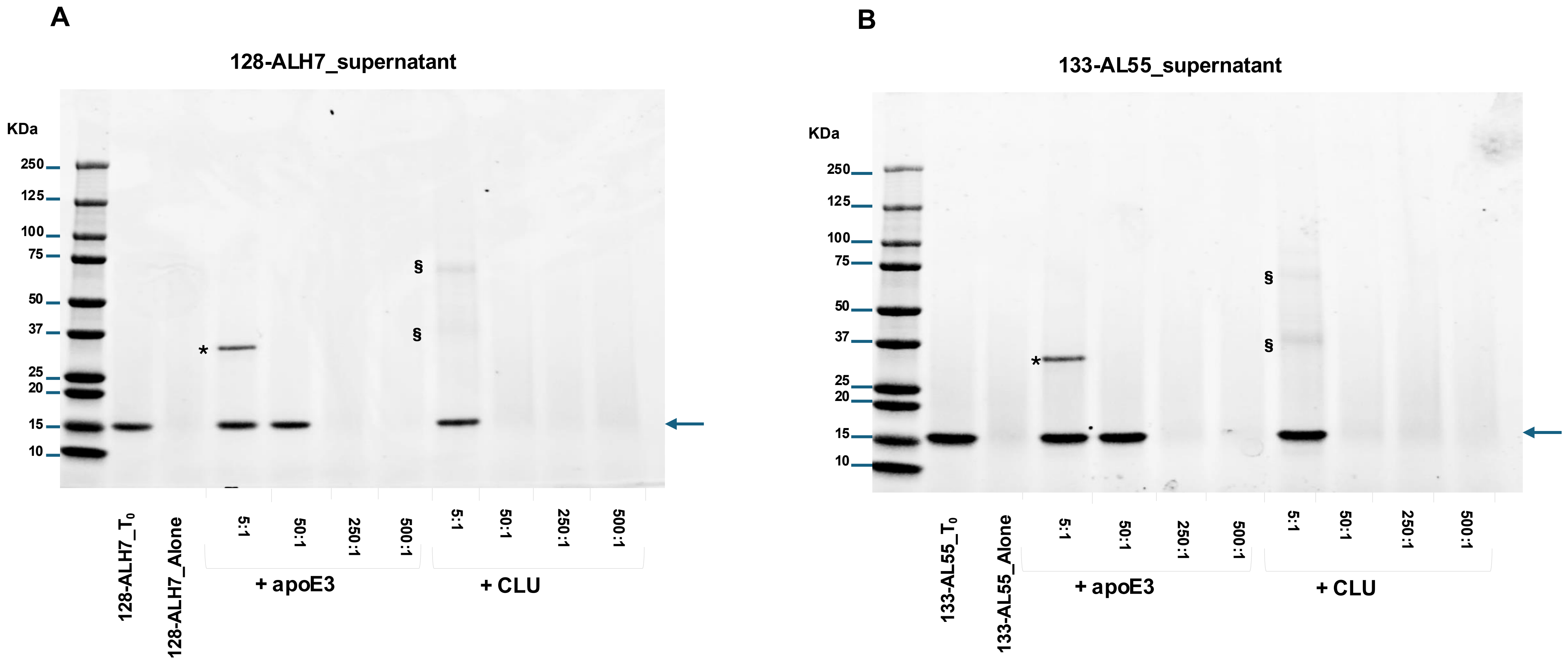
A
128-ALH7_supernatant
KDa
250
125
100
75
50
37
25
20
15
10
*
§
§
128-ALH7_T0
128-ALH7_Alone
5:1
50:1
250:1
500:1
5:1
50:1
250:1
500:1
+ apoE3
+ CLU
B
133-AL55_supernatant
KDa
250
125
100
75
50
37
25
20
15
10
*
§
§
133-AL55_T0
133-AL55_Alone
5:1
50:1
250:1
500:1
5:1
50:1
250:1
500:1
+ apoE3
+ CLU

Figure S3

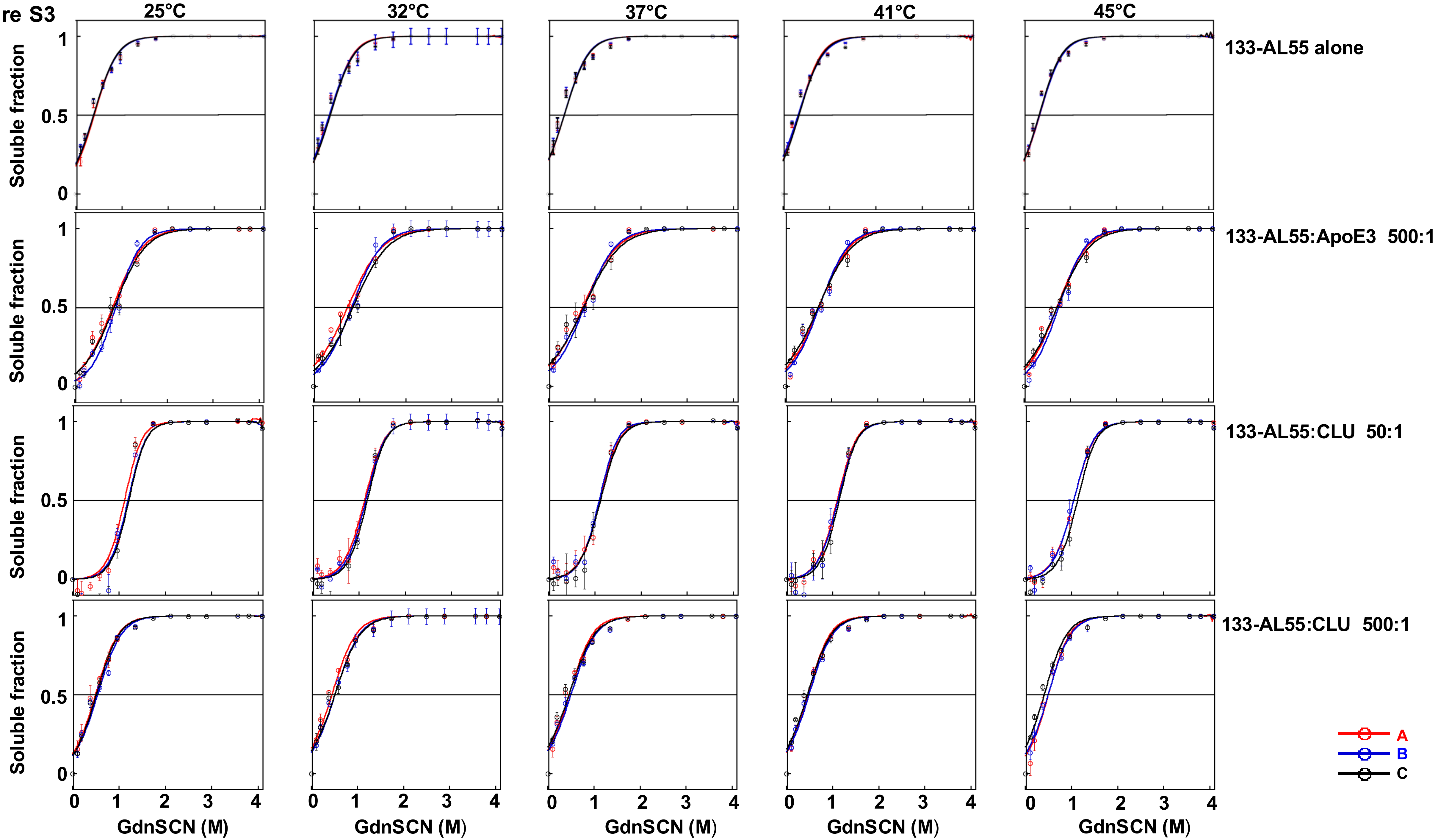
25°C
32°C
37°C
41°C
45°C
133-AL55 alone
133-AL55:ApoE3 500:1
133-AL55:CLU 50:1
133-AL55:CLU 500:1
Soluble fraction
0
0.5
1
GdnSCN (M)
0
1
2
3
4
A
B
C

Figure S4

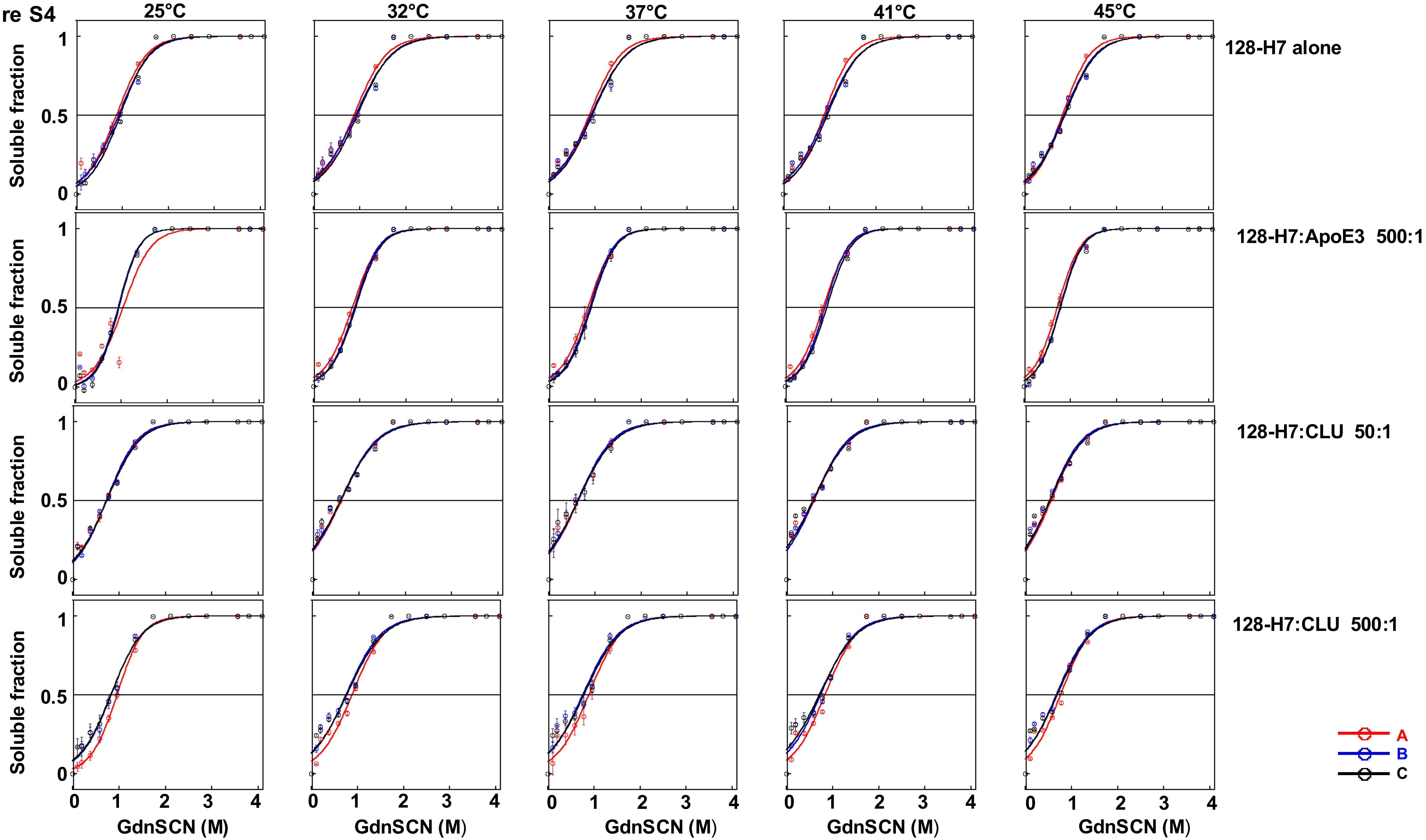
25°C
32°C
37°C
41°C
45°C
128-H7 alone
128-H7:ApoE3 500:1
128-H7:CLU 50:1
128-H7:CLU 500:1
Soluble fraction
GdnSCN (M)
A
B
C
